# HIV-1 Env gp41 Transmembrane Domain Dynamics are Modulated by Lipid, Water, and Ion Interactions

**DOI:** 10.1101/292326

**Authors:** L.R. Hollingsworth, J.A. Lemkul, D.R. Bevan, A.M. Brown

## Abstract

The gp41 transmembrane domain (TMD) of the envelope glycoprotein (Env) of the human immunodeficiency virus (HIV) modulates the conformation of the viral envelope spike, the only druggable target on the surface of the virion. Understanding of TMD dynamics is needed to better probe and target Env with small molecule and antibody therapies. However, little is known about TMD dynamics due to difficulties in describing native membrane properties. Here, we performed atomistic molecular dynamics simulations of a trimeric, prefusion TMD in a model, asymmetric viral membrane that mimics the native viral envelope. We found that water and chloride ions permeated the membrane and interacted with the highly conserved arginine bundle, (R696)_3_, at the center of the membrane and influenced TMD stability by creating a network of hydrogen bonds and electrostatic interactions. We propose that this (R696)_3_ - water - anion network plays an important role in viral fusion with the host cell by modulating protein conformational changes within the membrane. Additionally, R683 and R707 at the exofacial and cytofacial membrane-water interfaces, respectively, are anchored in the lipid headgroup region and serve as a junction point for stabilization of the termini. The membrane thins as a result of the tilting of the TMD trimer, with nearby lipids increasing in volume, leading to an entropic driving force for TMD conformational change. These results provide additional detail and perspective on the influence of certain lipid types on TMD dynamics and rationale for targeting key residues of the TMD for therapeutic design. These insights into the molecular details of TMD membrane anchoring will build towards a greater understanding of dynamics that lead to viral fusion with the host cell.

## Introduction

More than 36 million people worldwide currently live with human immunodeficiency virus (HIV), and a vaccine remains elusive (1). The sole antigenic target on HIV is the trimeric envelope glycoprotein (Env), which is comprised of heterodimeric subunits that include a surface glycoprotein (gp120) and a transmembrane glycoprotein (gp41) (2). Env mediates the entry of HIV into target cells through a cascade of conformational changes upon binding host cellular receptors (2). Briefly, Env first binds the CD4 receptor (3), leading to a global conformational shift of Env from the prefusion closed state to the activated open state, characterized by large rearrangements of the surface exposed glycoprotein regions in the ectodomain (4, 5). Binding of open state Env to either chemokine co-receptor, CCR5 or CXCR (4, 6, 7), leads to the formation of a six-helix transmembrane bundle that drives viral fusion with the host cell membrane (8, 9). Unbound Env samples the thermodynamic landscape between the prefusion closed state and the activated open state, hindering the ability to rationally design a broadly neutralizing vaccine (10–16). In addition to differences in dynamics due to mutations in the surface exposed ectodomain, conformational coupling of the gp41 transmembrane domain (TMD) to the ectodomain affects the dynamics and therefore antigenicity of Env (17, 18). Thus, elucidating the dynamics of the Env gp41 TMD may aid in engineering novel immunogens that elicit potent and broadly neutralizing antibodies (19–21).

The highly conserved gp41 TMD transmembrane domain (TMD) of Env (Figure 1) is anchored in a cholesterol-rich lipid bilayer (22) that is flanked by the membrane proximal external region (MPER) on the exofacial leaflet and the cytoplasmic tail (CT) on the cytofacial leaflet. In the prefusion state, either one (23) or three (21, 24, 25) single-pass α-helices span the membrane (Figure 1A). Previous studies have proposed that either oligomeric contacts (21, 24) or head group snorkeling (23) stabilizes an internal arginine, R696, which is buried in the membrane (21). Additionally, studies purport that a GxxxG motif in the TMD either stabilizes the trimer interface or acts as a signaling domain (26, 27). Despite this information and the importance of the TMD to HIV rational drug design, little is known about the dynamics of the TMD and its interactions within the HIV viral membrane, stimulating the present study.

**Figure 1.**
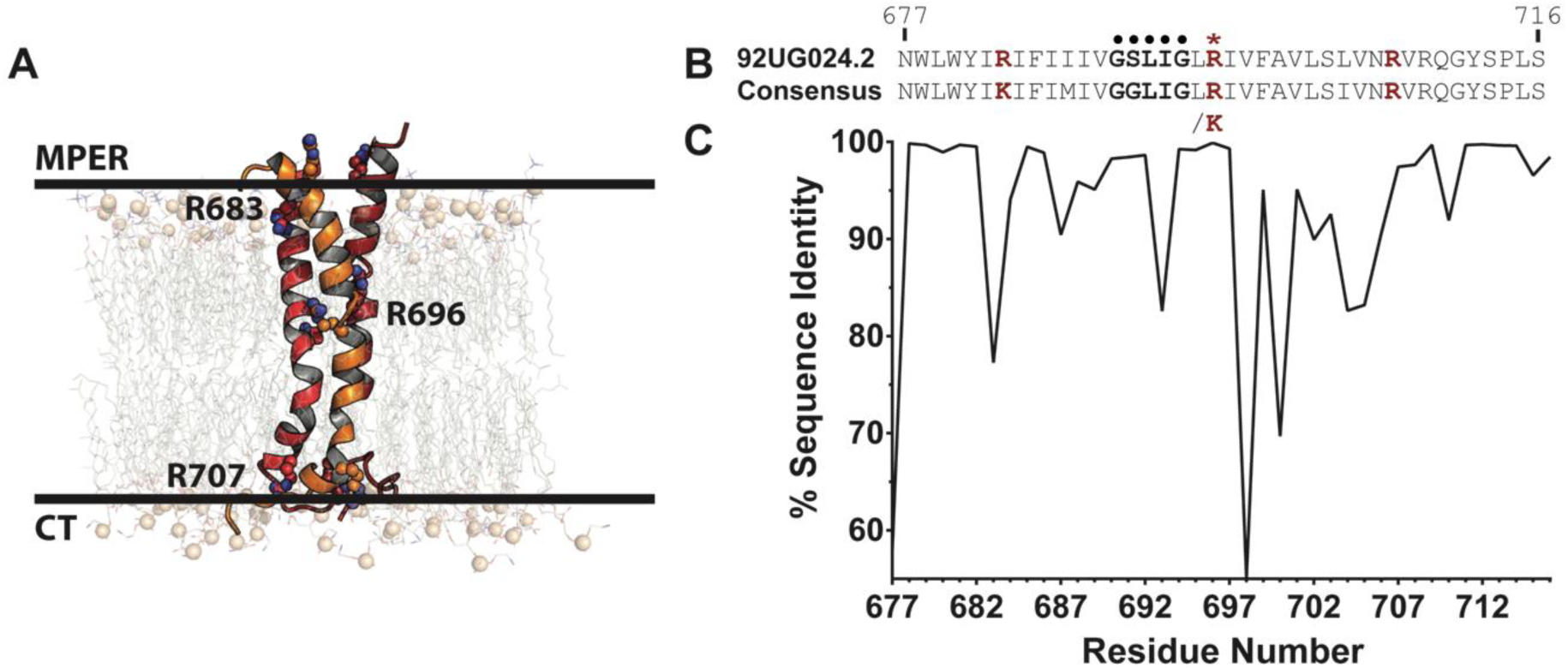
Properties of the TMD of gp41. (A) Representation of the HIV TMD embedded in an asymmetric membrane at the start of simulation (see Methods). The TMD trimer is shown as cartoon with each chain colored red, orange, or dark red. Key, conserved arginine residues are labeled and shown as spheres. Phosphorus atoms of lipid headgroups are shown as tan spheres for perspective. (B) Sequence comparison of TMD of the 92UG024.2 strain used in this study and (C) the consensus sequence of 6114 HIV-1 M group sequences. The GxxxG motif (•) and R696 (*) are highlighted, with all key arginine/lysine residues (R/K683, R/K696, and R707) shown in red.

All-atom molecular dynamics (MD) simulations offer a robust approach to reveal the dynamics of the TMD and its effect on membrane biophysical properties. However, previous MD simulations of the TMD were limited by the structural information that was available at the time of simulation, including factors such as the initial protein conformation, membrane composition, oligomeric state, R696 protonation state, and simulation duration (28–31). Prior TMD MD models were either derived by homology modeling using the HIV transmembrane viral protein U (Vpu) as template (28–30) or from a 20-amino acid transmembrane *de novo* model (31) rather than the recently published trimeric NMR TMD structure (21) which was used in this study. Previous simulations that used monomeric transmembrane models (28–30) lack quaternary contacts that are believed to stabilize the internal arginine (R696) (21, 24, 25). Furthermore, previous MD simulations were limited in sampling and membrane composition and were therefore unable to reveal the complex properties of the membrane components and their effect on TMD dynamics (28–31). More recently, discussion has formed around the ability of the TMD to exist in reconstituted bicelles as a monomer (23) and the potential role of key arginine residues in stabilizing either a monomeric or a trimeric transmembrane structure (24). The positioning, dynamics, and relationship between lipid type and protein conformational change are of current interest given these recent findings and can be further explored in more atomistic detail with MD simulations. Herein, we conducted MD simulations that closely represent the native viral membrane by explicitly modeling membrane lipid composition and asymmetry. We postulated that such an approach would provide novel information relevant to the atomistic and molecular dynamics that occur in this TMD, as influenced by the membrane environment, and serve as a basis for future simulations as more membrane associated structures are solved.

## Methods

### Sequence Analysis

Aligned envelope glycoprotein sequences from the HIV-1 M group were obtained from the Los Alamos HIV Database (http://www.hiv.lanl.gov/). Regions corresponding to the NMR TMD structure (from HIV-1 strain 92UG024.2 - PDB ID: 5JYN (21)) indices (database aligned indices 1242-1287) were parsed. The total dataset contained 6114 sequences. Sequence identity was analyzed with the Schrodinger Suite v2017-1 (32). We specifically focused on the HIV-1 virus, and the M group strain, which is responsible for the majority (M) of the global HIV epidemic (33).

### System Construction and MD Simulations

The cholesterol-containing phospholipid membrane system was created with the replacement method using the Membrane Builder module (34, 35) in the CHARMM-GUI interface (36, 37) to model the composition of the cholesterol-rich native HIV membrane (22) (Table 1). Due to the absence of plasmalogen lipid parameters in the CHARMM36 force field, a polyunsaturated fat, 1-stearoyl-2-docosahexaenoyl-sn-glycerophosphoethanolamine (SDPE), was used as a surrogate for plasmenylethanolamine (38). Prior work showed that there are similarities in simulation properties (density, thickness, deuterium order parameters) between a diacyl lipid and the corresponding plasmalogen (39). A 2.25-nm layer of water was placed on the exofacial and cytofacial leaflets of the lipid bilayer, resulting in a lipid hydration of approximately 46 waters per lipid.

**Table 1.**
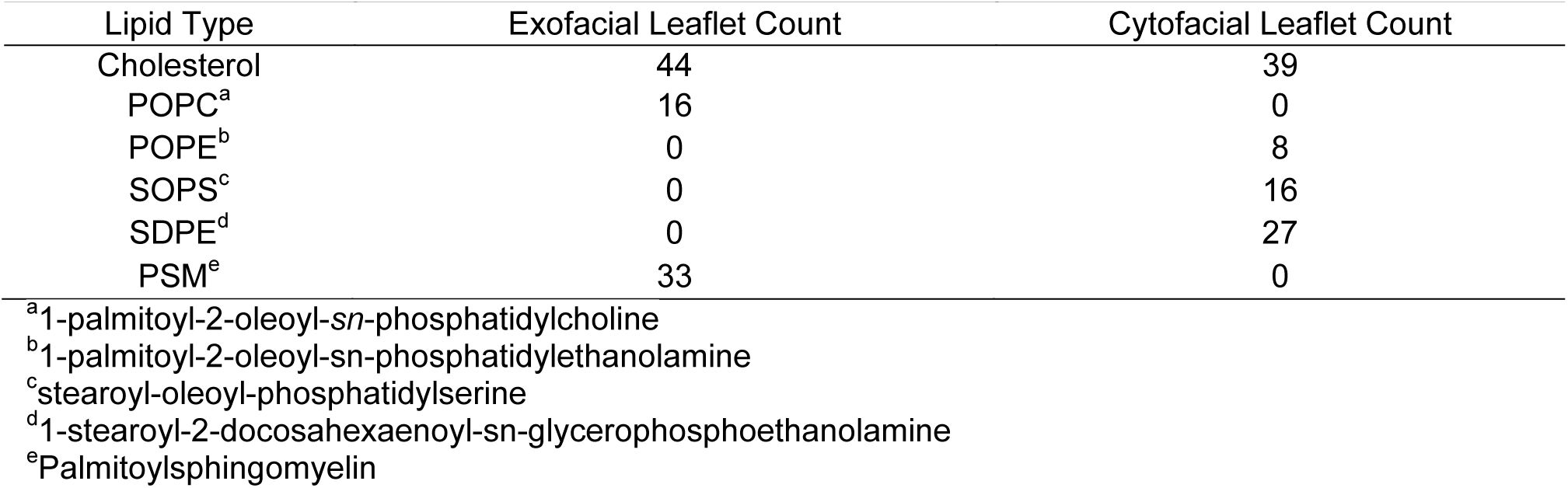
Membrane composition of the membrane-only simulation system.

| Lipid Type | Exofacial Leaflet Count | Cytofacial Leaflet Count |
| --- | --- | --- |
| Cholesterol | 44 | 39 |
| POPC<sup>a</sup> | 16 | 0 |
| POPE<sup>b</sup> | 0 | 8 |
| SOPS<sup>c</sup> | 0 | 16 |
| SDPE<sup>d</sup> | 0 | 27 |
| PSM<sup>e</sup> | 33 | 0 |

<sup>a</sup>1-palmitoyl-2-oleoyl-sn-phosphatidylcholine
<sup>b</sup>1-palmitoyl-2-oleoyl-sn-phosphatidylethanolamine
<sup>c</sup>stearoyl-oleoyl-phosphatidylserine
<sup>d</sup>1-stearoyl-2-docosahexaenoyl-sn-glycerophosphoethanolamine
<sup>e</sup>Palmitoylsphingomyelin

All MD simulations used the GROMACS software package, version 2016.3 (40, 41). The CHARMM36m force field was applied to the proteins and CHARMM36 parameters for lipids (42–44). The CHARMM-modified TIP3P (45–47) water model was used, and each system contained 150 mM NaCl, including neutralizing counterions. Ions were placed in the box randomly within the water. Standard CHARMM ion parameters were employed (48, 49).

Each system was energy-minimized using the steepest descent method. Following energy minimization, equilibration was performed using the standard Membrane Builder protocol (35, 37, 50), in which restraints on selected non-hydrogen atoms were slowly relaxed throughout equilibration steps. Briefly, the first phase of equilibration was carried out under an NVT ensemble for 50 ps with an integration time step of 1 fs using the Berendsen weak coupling method (51) to maintain temperature at 310 K. During this time, restraints on phospholipid P atoms and the cholesterol O3 atoms were applied with a force constant of 1000 kJ mol^-1^ nm^-2^. Following NVT equilibration, NPT equilibration was performed for a total of 325 ps, with the initial 25 ps carried out with a 1-fs time step and the remaining 300 ps with a 2-fs time step. During this phase, position restraints on the lipids were relaxed from 400 kJ mol^-1^ nm^-2^ to 0 kJ mol^-1^ nm^-2^. During NPT equilibration, pressure was maintained at 1 bar, semi-isotropically using the Berendsen weak coupling method (51). Following equilibration, production simulations were carried out in the absence of any restraints using the Nosé-Hoover (52, 53) thermostat and Parrinello-Rahman (53, 54) barostat to maintain temperature and pressure at 310 K and 1 bar, respectively. The fourth-order P-LINCS algorithm (55) was employed to constrain the lengths of bonds involving hydrogen atoms, allowing an integration time step of 2 fs. A buffered neighbor list was maintained with the Verlet method in GROMACS with a minimum cutoff of 1.2 nm for all nonbonded interactions. Electrostatic forces were calculated using the smooth particle mesh Ewald (PME) method (56, 57) using cubic interpolation, a Fourier grid spacing of 0.16 nm, and a real-space cutoff of 1.2 nm. Van der Waals forces were computed with the Lennard-Jones equation and smoothly switched to zero from 1.0 – 1.2 nm. Periodic boundary conditions were employed in all three spatial dimensions.

The membrane was simulated for 200 ns, after which time the trimeric NMR structure of the HIV-1 92UG024.2 strain gp 41 TMD (PDB ID: 5JYN (21)) was aligned along the membrane normal using LAMBADA (58). Lipids having any atom within 1.4 nm of the Cα atoms of the protein were removed using the InflateGRO method (59), modified in-house to accommodate multiple lipid types. The final membrane composition following protein insertion and lipid deletion is given in Table 2. Water molecules inside of the lipid bilayer were removed with an in-house script prior to minimization and equilibration using the protocol described above. For simulations containing the TMD trimer, position restraints were applied to all protein non-hydrogen atoms with a force constant of 1000 kJ mol^-1^ nm^-2^. The final contents and dimensions of all systems are shown in Table 3. Both systems (TMD-membrane and membrane-only) were simulated for 1.5 μs, and the TMD system was simulated in triplicate for a total of 4 production MD trajectories (three replicates of TMD-membrane trajectories totaling 4.5 μs and one control membrane-only trajectory totaling 1.5 μs). Area per lipid and bilayer thickness were calculated with GridMAT-MD (60) using a 200x200 grid for snapshots extracted at 1-ns intervals over the final 250 ns of simulation time. Tutorials and simulation input files can be found at our Open Science Framework page (https://osf.io/82n73/).

**Table 2.** Membrane composition of the transmembrane protein system following insertion of the gp41 trimer.

| Lipid Type | Total Number |
| --- | --- |
| Cholesterol | 80 |
| POPC<sup>a</sup> | 15 |
| POPE<sup>b</sup> | 8 |
| SOPS<sup>c</sup> | 16 |
| SDPE<sup>d</sup> | 25 |
| PSM<sup>e</sup> | 33 |

<sup>a</sup>1-palmitoyl-2-oleoyl-*sn*-phosphatidylcholine
<sup>b</sup>1-palmitoyl-2-oleoyl-*sn*-phosphatidylethanolamine
<sup>c</sup>stearoyl-oleoyl-phosphatidylserine
<sup>d</sup>1-stearoyl-2-docosahexaenoyl-*sn*-glycerophosphoethanolamine
<sup>e</sup>Palmitoylsphingomyelin

**Table 3.** MD simulation contents and box sizes of initial energy minimized systems prior to equilibration.

| System | # Cl<sup>-</sup> Ions | # Na<sup>+</sup> Ions | # Atoms | Box Side Lengths (X×Y×Z, nm) |
| --- | --- | --- | --- | --- |
| Membrane-only | 9 | 25 | 34854 | 6.20×6.20×8.49 |
| TMD (92UG024.2) | 30 | 34 | 34520 | 6.21×6.21×8.50 |

### MD Trajectory Analysis

Visual analysis and rendering was conducted with PyMOL (61), Chimera (62), and VMD (63). The final 250 ns from each trajectory, representing the equilibrated timeframe needed for membrane stabilization following TMD insertion (Figures S1-S2). Root-mean-square-deviation (RMSD) clustering was performed used the method of Daura *et al.* (64) with a 0.2-nm RMSD cutoff over the final 250 ns of each simulation (1.25-1.5 µs) to obtain representative protein conformations over the converged portion of each trajectory.

Trajectory data were analyzed with GROMACS (40, 41), CHARMM (60), and inhouse scripts. Hydrogen bonds between each TMD amino-acid sidechain and either water or lipid atoms were identified with the following criteria: 0.35-nm donor-acceptor heavy-atom distance and 30° hydrogen-donor-acceptor angular cutoff. Calculations were performed separately on each protein chain, and these data were combined to calculate averages and standard deviations across all three replicates.

To characterize the partitioning of water and ions into the membrane, the position of each water molecule or chloride ion relative to the TMD trimer was computed by calculating the z-coordinate of the center-of-mass (COM) of amino-acid Cα atoms at every fourth position along the membrane-spanning segment (residues 683-707). Any water oxygen atom or chloride ion that fell between the z-coordinate of the first and fourth residue of a given segment was recorded as occupying that vertical of the system.

Dynamics at the N- and C-termini were described by calculating the distances between each protein chain over time. The COM of residues R683 to I686 was used to describe the N-terminus, whereas the COM of residues L704 to R707 was used to describe the C-terminus. Each selection represents one full helical turn on the N- and C-terminal sides of the transmembrane domain. These distances were compiled into running averages over 1-ns blocks from snapshots saved every 10 ps.

To describe the distribution of lipids around each protein chain, radial distribution functions (RDF) were calculated relative to each protein chain independently. RDF were constructed for any lipid atom around any protein non-hydrogen atom.

Lipids were further characterized by their acyl chain tilt and molecular volume using CHARMM (60). To define tilt, a vector was constructed from the third carbon atom in the acyl chain to the carbon atom at the n-3 position. Doing so removes the influence of bending that occurs at the ester carbon atom (position 1 in the chain) and *gauche* conformational sampling that occurs at the terminal methyl group. The angle between this vector and the z-axis (corresponding to the membrane normal) was used to define acyl chain tilt. For each lipid, molecular volume was computed by constructing a Connelly surface around each lipid, with a water probe radius of 0.14 nm. The resulting volume enclosed by this surface was the molecular volume.

## Results and Discussion

To gain a greater understanding of the interplay between the HIV gp41 TMD and a viral membrane, we performed extensive MD simulations of the TMD trimer in a heterogeneous, asymmetric membrane. While it has been recently observed that a highly conserved arginine residue (R696) is a mediator of water exchange in the TMD trimer (24), we sought to further analyze the dynamics and movement of water, and potentially ions, into the trimer core. Additionally, we sought to provide further insight into the role of the highly conserved, positively charged residues that sit at the membrane-water interfaced based on the recent NMR structure (PDB ID: 5JYN). We specifically focused analysis on the dynamics of conserved TMD residues and the interactions of these residues with the membrane. By performing the longest simulations to date of the TMD trimer in a near-native viral membrane, we sought to resolve atomistic, biophysical questions related to the (i) influence of highly conserved, positively charged residues in the MPER (R683) and CT (R707) on water permeation and trimer stability, (ii) dynamic movements of the trimer complex relative to the flexible CT region, and (iii) effect and degree to which specific lipids contribute to trimer stability and conformational change.

### Dynamics of the TMD

The balance of interactions among the TMD, water, and membrane lipids has strong implications on the dynamics of the TMD trimer. To characterize these interactions, we computed the number of hydrogen bonds formed between each amino acid sidechain in the TMD with either water or each lipid type (Figure 2A,B and Figures S3 and S4). R683 is located at the interface of the exofacial leaflet and water, and consequently formed hydrogen bonds with both water molecules and lipid headgroup atoms. R683 consistently maintained a large number of hydrogen bonds with water molecules over the entire duration of the simulation (Figure 2A), whereas R683 participated in few hydrogen bonds with lipids over the final 250 ns (Figure 2B). We also observed that R683 sampled a very narrow range of positions along the z-axis (Figure 2C), indicating its preference to maintain water interactions over lipid contacts. Across all three individual R683 residue positions of each peptide in the trimer and in all three simulations, we observed that R683 formed approximately 9 ± 2 hydrogen bonds with water throughout the entire simulation (Figure 2A), supporting its role in being solvent-exposed and therefore accessible for binding antibodies and other therapeutics. Indeed, this interfacial positioning of R683 and its role in modulating interactions at the membrane interface is supported by studies of MPER-binding antibodies, such as 4E10 (65) and DH511.2 (66), which also bind to residues and lipids on the exofacial interface region. Hydrogen bonding of R683 sidechain to lipid headgroups in the exofacial leaflet increased from zero to two interactions each with both PSM and POPC over the course of the simulation, for a total of approximately 4 hydrogen bonds between all R683 residues of the trimer and lipids. This hydrogen bonding network at the exofacial interface suggests the combined effect of water and lipid hydrogen bonding to R683 maintains the position of the TMD trimer with binding to lipid headgroups as secondary. The large, positively charged sidechain of arginine is advantageous in this position, as the guanidinium moiety can simultaneously form hydrogen bonds to lipid phosphate groups and interfacial water molecules. To characterize the proximity of R683 to the membrane-water interface, we analyzed the distribution of guanidinium center-of-mass (COM) coordinates along the z-axis relative to exofacial lipid (POPC and PSM) P atoms. We found that the guanidinium moieties buried themselves ~0.1 nm below the phosphate groups (Figure S5). As a result of these interactions, positional fluctuations in the backbone at R683 were diminished relative to the flexible N-terminal residues (Figures 2D and S6).

**Figure 2.**
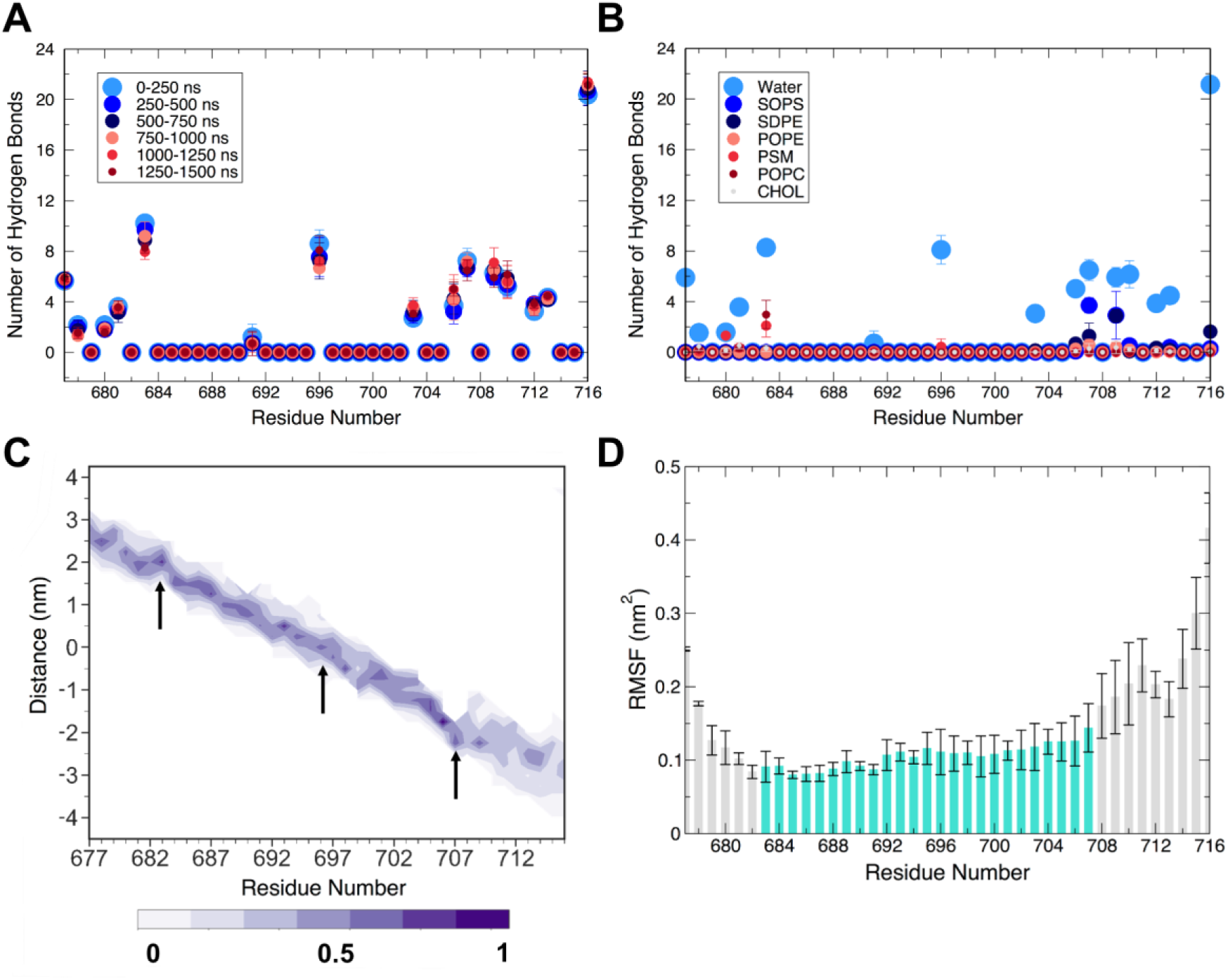
MD simulations elucidate membrane interface and protein dynamics of the TMD. (A) Average number of hydrogen bonds across all replicates of simulation between protein side chains and water molecules. (B) Average number of hydrogen bonds formed between protein side chains and water for each membrane lipid type over the last 250 ns of simulation. (C) Per-residue probability density (shaded bar) of z-coordinate over the last 250 ns of simulation relative to the membrane center of mass, showing membrane insertion. Important arginine residues (R683, R696, R707) are marked with arrows. (D) Average per-residue backbone RMSF over the final 250 ns of a simulation for all three replicates. The shaded region spans the membrane (683-707). Values and error bars shown in panels (A), (B), and (D) are the average and standard deviation, respectively, computed over the equivalent residues in the three replicates for the indicated time intervals.

While no mutations were assessed in the present study, Yi *et al.* (67) performed alanine mutagenesis throughout the MPER region (residues 660-683) in both a CXCR4 co-receptor utilizing strain (HXB2) and a CCR5 utilizing strain (JRFL) of HIV. Based on our sequence comparison, lysine is 80% conserved at position 683 across the 6114 sequences of the HIV-M group analyzed (Figure 1B), with an arginine residue at position 683 in the strain explored in this study (92UG024.2). Mutation of K683 to alanine in both the HXB2 and JRFL strain influenced viral entry, with either complete abolishment (HXB2) or diminished (JRFL) viral entry. This suggests that mutations at position 683 are influential to viral entry and are strain-specific, showing distinct sensitivity at this residue position Interestingly, a similar effect was observed at several preceding residues (L679, W680, and Y681) (67). These residues sit at the membrane-water interface and interacted less extensively with lipids, but did interact with water molecules (Figure 2A, B). Combined, these data support the hypothesis that positive charge is a key requirement at position 683 in the 92UG024.2 strain. Positive charge is necessary for solvent-accessibility and anchoring at the membrane-water interface, facilitating trimer positioning and HIV fusion. The interactions of R683 with water and lipid headgroups in our simulations show that a positively charged residue at this position forms a network of membrane-protein and membrane-water interactions to anchor the position of the TMD and promote conformational rigidity at the exofacial interface (Figures 2 and S6). The presence of a positively charged residue at position 683 could therefore be influential in modulating water at the extracellular membrane-water interface. Thus, conserved, charged residues in the MPER region may present a promising epitope for therapeutic design against HIV envelopes.

On the cytofacial face of the membrane, we observed a combined 8 ± 1 hydrogen bonds between water and the R707 residue of all three peptides (Figure 2A). These hydrogen bonds were preserved over the duration of the simulation, and increased hydrogen bonding with the headgroup of SOPS over time also was observed (from 1.0 ± 0.5 over 0-250 ns to 3 ± 1 over the last 250 ns). The advantageous positioning of R707 allows principal interactions with water and secondary interactions with the negatively charged headgroups of SOPS. This outcome is similar to our findings with R683, indicating that arginine serves a general role in forming networks of interactions among the protein, lipids, and interfacial waters. Additionally, R707 is positioned at the junction of the comparatively rigid TM region and more flexible C-terminal residues (Figure 2D), and its interactions with water and SOPS restrict its conformational freedom at the membrane-water interface (Figure 2C, arrow at position 707). Residues 708-716 sample greater z-coordinate space than 707, indicating that R707 maintains the structural positioning of the TMD trimer. We propose that R707 anchors into the lipid headgroup region to facilitate and stabilize the membrane-water interface, while allowing portions of the CT to fluctuate (Figures 2C and S6) to cause the passage of water and ions into the core of the structure (see below). We also observed that N706 forms some hydrogen bonds with water (4 ± 1), albeit fewer than R707 (8 ± 1, Figure 2A). Hydrogen bonding between N706 and PE-containing (SDPE, POPE) lipid headgroups was also observed to a lesser extent, with approximately 1 interaction being transiently observed over the duration of the simulation (Figure S4). Functional studies have shown that the N706A mutant has impaired fusion and infectivity, and an R707G mutant was fusion-deficient (21, 68). N706 and R707 are thus key in cytofacial leaflet anchoring and ultimately viral function. These data suggest that R707 plays a primary role in modulating protein-water interactions, with N706 acting as an additional contributor in positioning of the C-terminal region at the membrane-water interface.

We subsequently explored the conformational sampling of the three α-helices of the TMD relative to each other. Using the interfacial boundaries established previously, we measured the COM distance between MPER (towards the N-terminus of each peptide, Figure 1A) and CT residues (towards the C-terminus) of each pair of helices (Figure S7). The inter-helical distances at the MPER and CT sequences reflect the differing ability of these regions to open and allow the entry of water and ions into the trimer core (see below). The MPER region remains very stable and tightly packed with stable COM distances between neighboring peptides (~1.2 nm), consistent with experimental observations of the known α-helical character of MPER peptides (65), residue-specific NMR relaxation rates (24), and the observed flexibility of CT residues 707-751 (69). Minor increases in COM distance were observed in the MPER region, persisting for at most 100 ns before returning to a tightly packed state (Figure S7). Additionally, the MPER residues of the TMD manifested limited diffusion along the z-axis, as noted above for R683 (Figure 2C), suggesting a more packed, rigid helical bundle at the exofacial membrane interface.

In contrast, residues in the CT region exhibited rapid opening, which was maintained for long periods of time in each simulation (Figure S7). Typical increases in COM distance were on the order of 0.5 - 0.8 nm, typically involving one helix moving relative to the other helices. Opening of the trimeric bundle was a result of kinking of the CT half of the TMD (residues 695-707), (Figure S8). Fluctuations in the kink angles relative to the membrane normal (z-axis) corresponded to opening and closing events between CT residues of neighboring peptides (Figures S7 and S8). Simultaneous increase in kink angle between two neighboring peptides resulted in increased COM distance in these pairs, while opposing kink (e.g. one peptide straightened while the other kinked) resulted in no change in COM distance (Figure S8). Thus, the previously noted fluctuation at the C-terminal end of the TMD results from bending of the α-helices relative to the membrane normal.

The agreement of these MD simulations with experimental observations validates our approach and extends these observations by providing a molecular basis for the key interfacial stabilization roles of arginine residues R683 and R707. The above data rationalize the positioning of key arginine residues at the membrane-solvent interface, as well as providing evidence of differences in the dynamics of the two termini, ultimately connecting trimer structural stabilization with the ability to facilitate water and chloride permeation into the TMD

### Water and ion permeation into the TMD trimer

Arginine residues at position 696 are highly conserved (Figure 1B) and located at the bilayer core, motivating the present study. Strikingly, water molecules quickly partitioned into the trimer core to form hydrogen bonds with all three R696 residues (Figure 2A). Hydrogen bonding between water and any other residue in the TMD was negligible, and R696 did not participate in hydrogen bonding interactions with lipids (Figure 2B). Based on this observation, which requires water molecules to enter the membrane, water molecules and chloride ions were assessed for their potential to penetrate the lipid bilayer (Figure 3). The membrane-spanning region was defined in terms of the protein, with cross-sections defined as the z-coordinate of the Cα atoms of four consecutive residues, spanning the entire TM region (residues 683-707). The z-coordinate of each water oxygen atom or chloride atom were recorded, and if they fell between the maximum and minimum z-coordinate of the chosen four-residue segment, they were counted as occupying this membrane-spanning region. In this way, it was possible to track the accumulation of water in the TM region. Results of water and chloride ion counting are shown in Figure 3A,B. A large number of water molecules were observed at z positions within R683-I687 and S703-R707, though these include water molecules at the membrane-water interface, thus not bound within the TMD helix bundle. Water molecules accumulated within the trimer core (Figure 3A), though these water networks were not continuous throughout the trimer core (Figure 3C). Amino-acid residues in the middle of the sequence disrupted the water network. These findings are consistent with recent hydrogen-deuterium exchange experiments, wherein D_2_O permeated through the cytofacial side of the TMD but was blocked by the hydrophilic core above R696 (24). These findings were consistent across the three replicate simulations (Figure S9).

**Figure 3.**
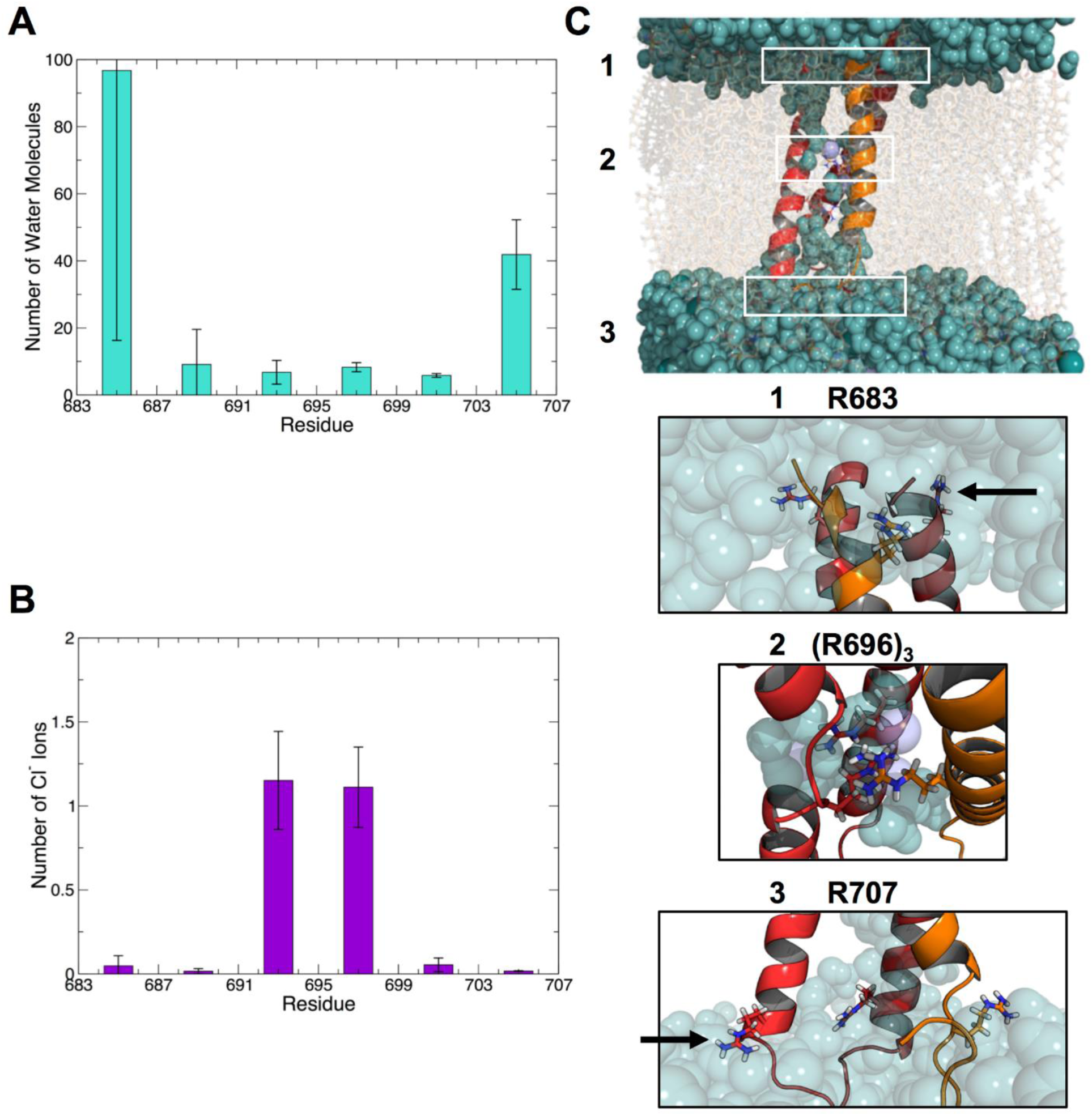
Water and chloride ions permeate the membrane to stabilize the three-helix bundle. Average number of (A) water molecules and (B) chloride ions between the indicated residues over the last 250 ns of simulation time. Bars indicate the average number of water molecules or chloride ions between the intervals of four residues (labeled on the x-axis). Averages and standard deviations were computed over each of the three replicate simulations. (C) Cartoon of a dominant morphology (48.6% of frames) from a representative replicate. Water molecules (teal) and chloride ions (purple) are shown as spheres. Expanded views of arginine residues in the center of the membrane (R696) and at the membrane-water interfaces (R683 and R707) are shown for reference. The indicated arginine residues are shown in stick.

Similarly, chloride ions diffused into the hydrated protein trimer core. On average, two ions were bound in the vicinity of R696 (Figure 3B,C). These ions tended to associate with R696, thereby offsetting the concentrated positive charge that arises from the proximity of the three R696 residues. It is notable that the chloride ions had a strong preference for this environment, as few, if any, chloride ions were observed at any other position within the TM region (Figure 3B). Water and ions z-coordinate position sampling over the last 250 ns of each simulation are shown in Figures S10 and S11.

Previous MD studies showed that buried arginine and lysine residues, like R696, often facilitate the formation of such water networks; however, arginine residues can form more extensive water networks than lysine residues due to their chemical structure, as the guanidinium moiety can participate in a greater number of simultaneous hydrogen bonding interactions (70, 71). Across HIV-1 M group strains, R696 is 98.2% conserved (6005 sequences), whereas K696 is observed in 1.68% percent of strains (103 sequences, Figure 1C). Thus, a positively charged residue is almost completely conserved at this residue positions. Only three HIV strains used in the conservation scoring do not have a positive residue at position 696, as strains QH0705 (AF277066) (72), R214 (AY773339) (73), and 10NG040248 (KX389608) (74) have serine, cysteine, and glycine substitutions, respectively. Thus, the dominant presence of a positively charged residue in this position suggests that a residue at the core of the bilayer facilitates a network of water-residue hydrogen bonding that likely plays a critical functional role in stabilizing the α-helical bundle, modulating the stability of the trimer and supporting the trimer integrity in an asymmetric membrane environment that is typical for viral fusion.

Several mutagenesis studies have been performed on R696, with R696A reducing cell-cell fusion and R696L eliminating fusion (21, 68). Given the conservation of charged residues at position 696 and the observation of water and chloride ion interactions at this interface, the combination of previous experiments showing water exchange into the trimer core (24) and the simulations performed here suggests that water serves to screen the high charge density at the trimer interface and facilitates the formation of the network of interactions formed by R696 to stabilize the trimer unit that are required for subsequent membrane-fusion events.

### Effects of lipid asymmetry on membrane properties and protein dynamics

In addition to membrane hydration, the presence of the TMD perturbed the membrane structure (Figure 4). The lateral area per lipid (APL) of both leaflets from each independent replicate of the TMD-membrane simulations was significantly reduced compared to the membrane-only control (Figure 4A). Further, the thickness of the membrane decreased, particularly in local deformations around the transmembrane helices (Figure 4B). NMR and MD studies of other molecular systems showed that arginine and membrane-flanking residues can cause membrane thinning to accommodate stabilization of hydrophilic or charged sidechains (75, 76). Taken together, the observed membrane thinning is likely due to the interfacial interactions of R683 and R707 on the exofacial and cytofacial leaflets, respectively (Figure 3C).

**Figure 4.**
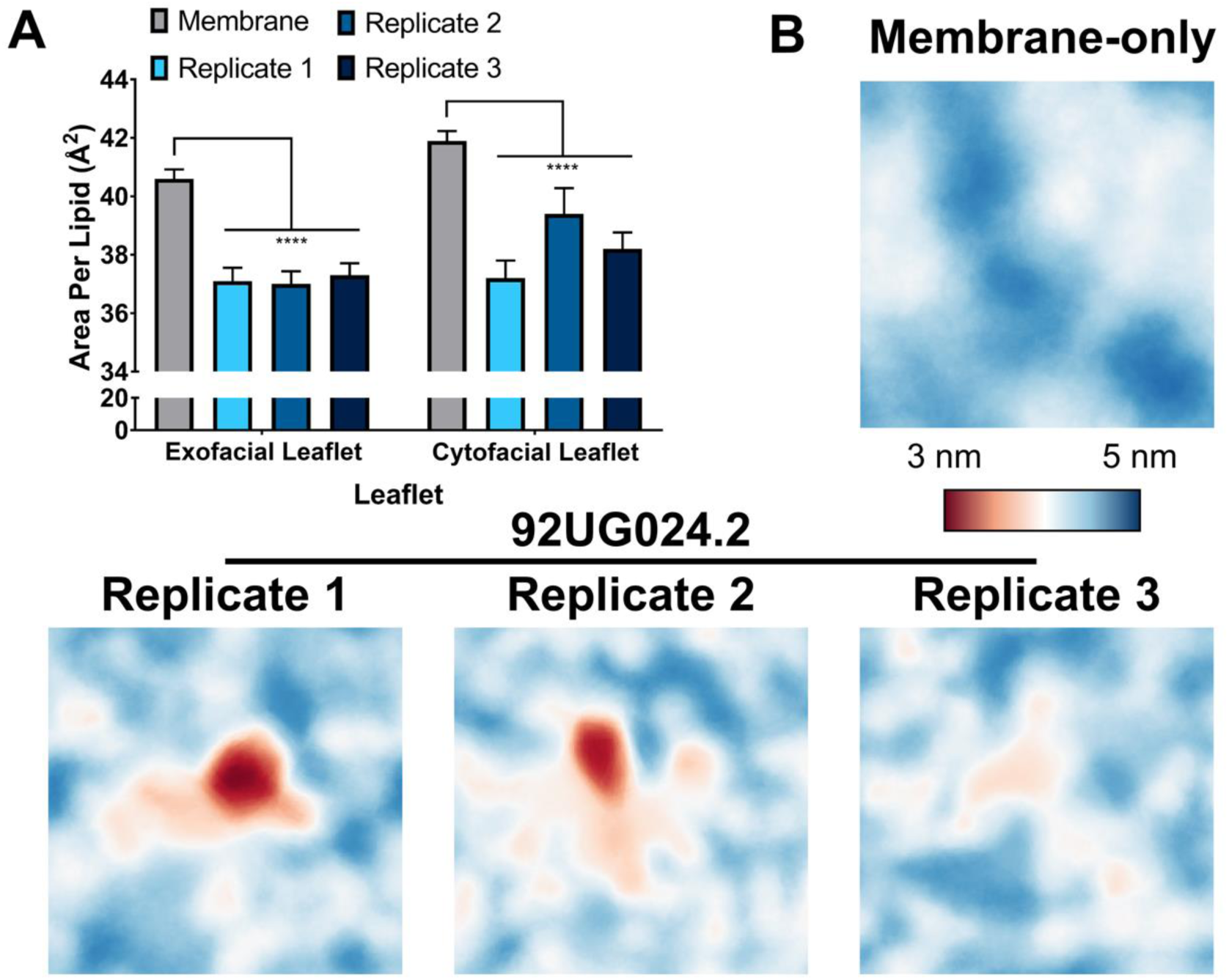
Transmembrane gp41 induces membrane thinning and perturbs membrane thickness. (A) Average area per lipid over the final 250 ns of each simulation. Error bar values represent ± standard deviation (n = 2501, Two-way ANOVA with Sidak post-test, ****p < 0.0001). (B) Average membrane thicknesses over the final 250 ns of each simulation. Large local membrane perturbations occur in the presence of transmembrane gp41.

Phospholipid composition and membrane properties have been observed to affect water ordering at membrane interfaces (77). Thus, the HIV membrane lipid composition (22, 78) and local perturbations in lipid properties (Figure 4) likely contribute to both viral entry and the potency and/or specificity of antibodies targeting the MPER. These properties are inherently a function of both protein and lipid dynamics. To describe the distribution of lipids around the TMD and identify any specific interactions of importance, we computed radial distribution functions (RDF) of lipid occupancy around each protein chain of the trimer. On the exofacial leaflet, POPC, PSM, and CHOL showed no preference for interacting with the TMD (Figures S12 and S13). In contrast, in the cytofacial leaflet, POPE and SDPE were modestly enriched around the protein over time, with POPE approaching the TMD trimer more closely and SDPE accumulating at a radial distance of ~1 nm (Figure S13, S14). The remaining lipids in the cytofacial leaflet (SOPS and CHOL) were distributed homogeneously.

The molecular volume of a lipid is related to APL and serves as an additional descriptor of the packing within the membrane. We computed the molecular volumes of POPE and SDPE lipids, finding that there was a bimodal distribution for each (Figure 5A). The smaller peak at larger volume led us to investigate whether or not there was an impact of the protein on the lipid volumes. To this end, we plotted the individual lipid molecular volumes as a function of the minimum distance between the lipid and any peptide chain, using only non-hydrogen atoms for the calculation. Probabilities of lipid volumes as a function of distance were converted to free energy (Figure 5B,C), G = - kTln[P(V_m_,r)], where P is the probability of a given volume (V_m_) occurring at a given distance (r). Results of these calculations are shown in Figure 5B,C. As the lipids packed against the TMD, their molecular volumes increased. This behavior suggests an entropic driving force for this association. The peptide chains sample a wide range of tilt angles, with dominant sampling around 5°, 16°, and 28° (Figure S15). The average peptide tilt angle over the last 250 ns of all simulations was 16.7° ± 7.5°, and the unsaturated chains of POPE and SDPE mirrored this value. In SDPE, the docosahexaenoyl chain had an average tilt of 16.7° ± 11.7° and in POPE, the oleoyl chain had an average tilt of 16.3° ± 10.6°.

**Figure 5.**
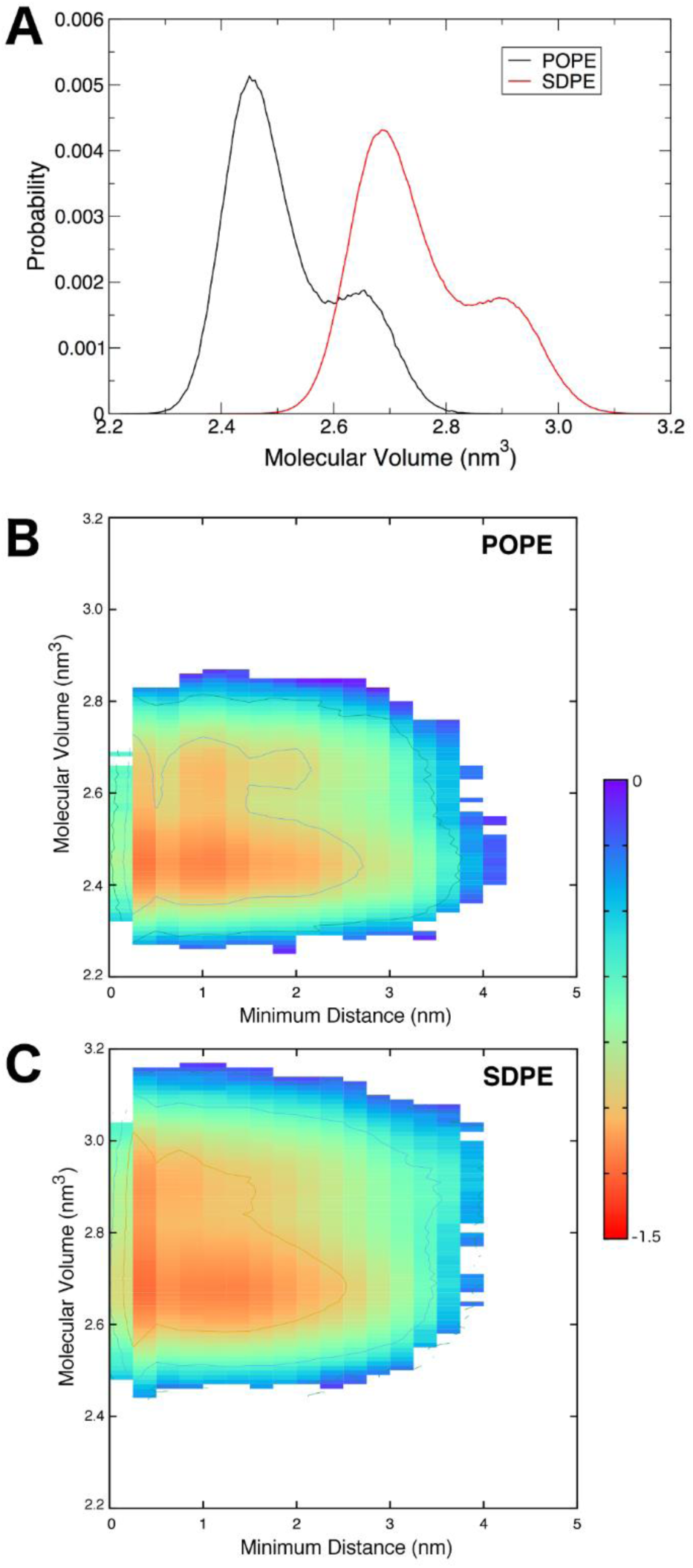
Dynamics of membrane lipids. (A) Molecular volume distributions of POPE and SDPE lipids in the cytofacial leaflet. (B,C) Molecular volumes of POPE and SDPE as a function of their minimum distance to the gp41 trimer, plotted as free energy surfaces. The free energy scale (in kcal mol^-1^) on the right applies to both panels (B) and (C).

The complementarity of the lipid geometries with the peptides suggests that unsaturated lipids are particularly amenable to the tilt in the TMD trimer that is required for it to take up water and ions as part of its shift in conformational ensemble. Together, these data suggest an entropic driving force for the conformational changes observed in the TMD. That is, tilting of the transmembrane helices allows the trimer to sample a greater volume, as the structure converts from sampling a cylindrical volume to a conical volume. This process is driven by helical precession entropy, which is used to overcome the free energy barrier to membrane deformation and contributes to the membrane thinning shown in Figure 4 (79). The commensurate increase in lipid molecular volume upon approaching the trimer gives rise to a similar entropic gain in the lipids. Thus, the interactions of the TMD with the asymmetric membrane are characterized by hydrogen bonding interactions, contributing to favorable enthalpy, as well as entropic driving forces that promote conformational change as a function of specific protein-lipid interactions.

## Conclusions

We have created a dynamic model for the gp41 TMD of the HIV-1 Env in a lipid bilayer and show that results from MD simulations are consistent with available biophysical and cellular experimental data. Additionally, by using an asymmetric membrane that contains near-native viral lipid composition, we have advanced towards understanding the role of conserved residues, protein dynamics, and sidechain positioning that influence the dynamics and membrane interactions of the prefusion TMD trimer. Through these simulations, we have obtained a greater structural and thermodynamic characterization of this important viral protein. Our data suggest that water molecules and anions permeate transmembrane Env to stabilize the buried R696 trimer and that the membrane likely thins as a result of peptide tilting and subsequent increases in nearby lipid molecular volumes, suggesting an entropic driving force for conformational change and membrane perturbation. These results provide a basis for rational therapeutic design targeting the transmembrane region and MPER and will be increasingly powerful as the relationship between the antigenic surface of Env and the transmembrane domain becomes apparent.

## Acknowledgments

The authors thank Advanced Research Computing at Virginia Tech for high performance computing resources and Priya Veeraraghavan for technical advice and for critical reading of the manuscript.

## Author Contributions

L.R.H, A.M.B, and D.R.B. designed research protocol. L.R.H. performed simulations. A.M.B, J.A.L. and L.R.H. analyzed data. A.M.B. J.A.L., L.R.H, and D.R.B. wrote the paper.

## References

1. WHO HIV/AIDS Data. World Health Organization.

2. Wyatt, R., and J. Sodroski. 1998. The HIV-1 Envelope Glycoproteins: Fusogens, Antigens, and Immunogens. Science 280:1884–1888.

3. Dalgleish, A. G., P. C. L. Beverley, P. R. Clapham, D. H. Crawford, M. F. Greaves, and R. A. Weiss. 1984. The CD4 (T4) antigen is an essential component of the receptor for the AIDS retrovirus. Nature 312:763–767.

4. Ozorowski, G., J. Pallesen, N. de Val, D. Lyumkis, C. A. Cottrell, J. L. Torres, J. Copps, R. L. Stanfield, A. Cupo, P. Pugach, J. P. Moore, I. A. Wilson, and A. B. Ward. 2017. Open and closed structures reveal allostery and pliability in the HIV-1 envelope spike. Nature.

5. Liu, J., A. Bartesaghi, M. J. Borgnia, G. Sapiro, and S. Subramaniam. 2008. Molecular architecture of native HIV-1 gp120 trimers. Nature 455:109–113.

6. Choe, H., M. Farzan, Y. Sun, N. Sullivan, B. Rollins, P. D. Ponath, L. Wu, C. R. Mackay, G. LaRosa, W. Newman, N. Gerard, C. Gerard, and J. Sodroski. 1996. The beta-chemokine receptors CCR3 and CCR5 facilitate infection by primary HIV-1 isolates. Cell 85:1135–1148.

7. Feng, Y., C. C. Broder, P. E. Kennedy, and E. A. Berger. 1996. HIV-1 entry cofactor: functional cDNA cloning of a seven-transmembrane, G protein-coupled receptor. Science 272:872–877.

8. Chan, D. C., D. Fass, J. M. Berger, and P. S. Kim. 1997. Core Structure of gp41 from the HIV Envelope Glycoprotein. Cell 89:263–273.

9. Markosyan, R. M., M. Y. Leung, and F. S. Cohen. 2009. The Six-Helix Bundle of Human Immunodeficiency Virus Env Controls Pore Formation and Enlargement and Is Initiated at Residues Proximal to the Hairpin Turn. J. Virol. 83:10048–10057.

10. Guttman, M., A. Cupo, J.-P. Julien, R. W. Sanders, I. A. Wilson, J. P. Moore, and K. K. Lee. 2015. Antibody potency relates to the ability to recognize the closed, pre-fusion form of HIV Env. Nat. Commun. 6.

11. Munro, J. B., J. Gorman, X. Ma, Z. Zhou, J. Arthos, D. R. Burton, W. C. Koff, J. R. Courter, A. B. Smith, P. D. Kwong, S. C. Blanchard, and W. Mothes. 2014. Conformational dynamics of single HIV-1 envelope trimers on the surface of native virions. Science 346:759–763.

12. Herschhorn, A., X. Ma, C. Gu, J. D. Ventura, L. Castillo-Menendez, B. Melillo, D. S. Terry, A. B. Smith, S. C. Blanchard, J. B. Munro, W. Mothes, A. Finzi, and J. Sodroski. 2016. Release of gp120 Restraints Leads to an Entry-Competent Intermediate State of the HIV-1 Envelope Glycoproteins. mBio 7.

13. Pancera, M., T. Zhou, A. Druz, I. S. Georgiev, C. Soto, J. Gorman, J. Huang, P. Acharya, G. Y. Chuang, G. Ofek, G. B. Stewart-Jones, J. Stuckey, R. T. Bailer, M. G. Joyce, M. K. Louder, N. Tumba, Y. Yang, B. Zhang, M. S. Cohen, B. F. Haynes, J. R. Mascola, L. Morris, J. B. Munro, S. C. Blanchard, W. Mothes, M. Connors, and P. D. Kwong. 2014. Structure and immune recognition of trimeric pre-fusion HIV-1 Env. Nature 514:455–461.

14. Davenport, T. M., M. Guttman, W. Guo, B. Cleveland, M. Kahn, S.-L. Hu, and K. K. Lee. 2013. Isolate-Specific Differences in the Conformational Dynamics and Antigenicity of HIV-1 gp120. J. Virol.

15. Cai, Y., S. Karaca-Griffin, J. Chen, S. Tian, N. Fredette, C. E. Linton, S. Rits-Volloch, J. Lu, K. Wagh, J. Theiler, B. Korber, M. S. Seaman, S. C. Harrison, A. Carfi, and B. Chen. 2017. Antigenicity-defined conformations of an extremely neutralization-resistant HIV-1 envelope spike. Proc. Natl. Acad. Sci. U. S. A. 114:4477–4482.

16. Ward, A. B., and I. A. Wilson. 2017. The HIV-1 envelope glycoprotein structure: nailing down a moving target. Immunol. Rev. 275:21–32.

17. Hollingsworth, L. R. I. V., A. M. Brown, R. D. Gandour, and D. R. Bevan. 2018. Computational study of HIV gp120 as a target for polyanionic entry inhibitors: Exploiting the V3 loop region. PLoS One 13:e0190658.

18. Yokoyama, M., S. Naganawa, K. Yoshimura, S. Matsushita, and H. Sato. 2012. Structural Dynamics of HIV-1 Envelope Gp120 Outer Domain with V3 Loop. PLoS One 7:e37530.

19. Kondo, N., K. Miyauchi, F. Meng, A. Iwamoto, and Z. Matsuda. 2010. Conformational Changes of the HIV-1 Envelope Protein during Membrane Fusion Are Inhibited by the Replacement of Its Membrane-spanning Domain. J. Biol. Chem. 285:14681–14688.

20. Chen, J., J. M. Kovacs, H. Peng, S. Rits-Volloch, J. Lu, D. Park, E. Zablowsky, M. S. Seaman, and B. Chen. 2015. Effect of the cytoplasmic domain on antigenic characteristics of HIV-1 envelope glycoprotein. Science 349:191.

21. Dev, J., D. Park, Q. Fu, J. Chen, H. J. Ha, F. Ghantous, T. Herrmann, W. Chang, Z. Liu, G. Frey, M. S. Seaman, B. Chen, and J. J. Chou. 2016. Structural basis for membrane anchoring of HIV-1 envelope spike. Science 353:172.

22. Brügger, B., B. Glass, P. Haberkant, I. Leibrecht, F. T. Wieland, and H.-G. Kräusslich. 2006. The HIV lipidome: A raft with an unusual composition. Proc. Natl. Acad. Sci. U. S. A. 103:2641–2646.

23. Chiliveri, S. C., J. M. Louis, R. Ghirlando, J. L. Baber, and A. Bax. 2018. Tilted, Uninterrupted, Monomeric HIV-1 gp41 Transmembrane Helix from Residual Dipolar Couplings. Journal of the American Chemical Society 140:34–37.

24. Piai, A., J. Dev, Q. Fu, and J. J. Chou. 2017. Stability and Water Accessibility of the Trimeric Membrane Anchors of the HIV-1 Envelope Spikes. Journal of the American Chemical Society 139:18432–18435.

25. Reichart, T. M., M. M. Baksh, J.-K. Rhee, J. D. Fiedler, S. G. Sligar, M. G. Finn, M. B. Zwick, and P. E. Dawson. 2016. Trimerization of the HIV Transmembrane Domain in Lipid Bilayers Modulates Broadly Neutralizing Antibody Binding. Angew. Chem. 55:2688–2692.

26. Miyauchi, K., A. R. Curran, Y. Long, N. Kondo, A. Iwamoto, D. M. Engelman, and Z. Matsuda. 2010. The membrane-spanning domain of gp41 plays a critical role in intracellular trafficking of the HIV envelope protein. Retrovirology 7:95.

27. Teese, M. G., and D. Langosch. 2015. Role of GxxxG Motifs in Transmembrane Domain Interactions. Biochemistry 54:5125–5135.

28. Gangupomu, V. K., and C. F. Abrams. All-Atom Models of the Membrane-Spanning Domain of HIV-1 gp41 from Metadynamics. Biophys. J. 99:3438–3444.

29. Baker, M. K., and C. F. Abrams. 2014. Dynamics of Lipids, Cholesterol, and Transmembrane α-Helices from Microsecond Molecular Dynamics Simulations. J. Phys. Chem. B 118:13590–13600.

30. Baker, M. K., V. K. Gangupomu, and C. F. Abrams. 2014. Characterization of the water defect at the HIV-1 gp41 membrane spanning domain in bilayers with and without cholesterol using molecular simulations. Biochim. Biophys. Acta 1838:1396–1405.

31. Kim, J. H., T. L. Hartley, A. R. Curran, and D. M. Engelman. 2009. Molecular dynamics studies of the transmembrane domain of gp41 from HIV-1. Biochim. Biophys. Acta 1788:1804–1812.

32. 2017. Schrödinger Release 2017-1. Schrödinger, LLC, New York, NY.

33. Buonaguro, L., M. L. Tornesello, and F. M. Buonaguro. 2007. Human Immunodeficiency Virus Type 1 Subtype Distribution in the Worldwide Epidemic, Pathogenetic and Therapeutic Implications. Journal of Virology 81:10209–10219.

34. Jo, S., J. B. Lim, J. B. Klauda, and W. Im. 2009. CHARMM-GUI Membrane Builder for Mixed Bilayers and Its Application to Yeast Membranes. Biophys. J. 97:50–58.

35. Wu, E. L., X. Cheng, S. Jo, H. Rui, K. C. Song, E. M. Dávila-Contreras, Y. Qi, J. Lee, V. Monje-Galvan, R. M. Venable, J. B. Klauda, and W. Im. 2014. CHARMM-GUI Membrane Builder toward realistic biological membrane simulations. J. Comput. Chem. 35:1997–2004.

36. Jo, S., T. Kim, V. G. Iyer, and W. Im. 2008. CHARMM-GUI: A web-based graphical user interface for CHARMM. J. Comput. Chem. 29:1859–1865.

37. Lee, J., X. Cheng, J. M. Swails, M. S. Yeom, P. K. Eastman, J. A. Lemkul, S. Wei, J. Buckner, J. C. Jeong, Y. Qi, S. Jo, V. S. Pande, D. A. Case, C. L. Brooks, A. D. MacKerell, J. B. Klauda, and W. Im. 2016. CHARMM-GUI Input Generator for NAMD, GROMACS, AMBER, OpenMM, and CHARMM/OpenMM Simulations Using the CHARMM36 Additive Force Field. J. Chem. Theory Comput. 12:405–413.

38. Klauda, J. B., V. Monje, T. Kim, and W. Im. 2012. Improving the CHARMM Force Field for Polyunsaturated Fatty Acid Chains. J. Phys. Chem. B 116:9424–9431.

39. Rog, T., and A. Koivuniemi. 2016. The biophysical properties of ethanolamine plasmalogens revealed by atomistic molecular dynamics simulations. Biochim. Biophys. Acta 1858:97–103.

40. Páll, S., M. J. Abraham, C. Kutzner, B. Hess, and E. Lindahl. 2015. Tackling Exascale Software Challenges in Molecular Dynamics Simulations with GROMACS. In Solving Software Challenges for Exascale: International Conference on Exascale Applications and Software, EASC 2014, Stockholm, Sweden, April 2-3, 2014, Revised Selected Papers. S. Markidis, and E. Laure, editors. Springer International Publishing, Cham. 3–27.

41. Abraham, M. J., T. Murtola, R. Schulz, S. Páll, J. C. Smith, B. Hess, and E. Lindahl. 2015. GROMACS: High performance molecular simulations through multi-level parallelism from laptops to supercomputers. SoftwareX 1–2:19–25.

42. Huang, J., S. Rauscher, G. Nawrocki, T. Ran, M. Feig, B. L. de Groot, H. Grubmuller, and A. D. MacKerell Jr. 2017. CHARMM36m: an improved force field for folded and intrinsically disordered proteins. Nat. Meth. 14:71–73.

43. Venable, Richard M., Alexander J. Sodt, B. Rogaski, H. Rui, E. Hatcher, Alexander D. MacKerell Jr, Richard W. Pastor, and Jeffery B. Klauda. 2014. CHARMM All-Atom Additive Force Field for Sphingomyelin: Elucidation of Hydrogen Bonding and of Positive Curvature. Biophys. J. 107:134–145.

44. Klauda, J. B., R. M. Venable, J. A. Freites, J. W. O’Connor, D. J. Tobias, C. Mondragon-Ramirez, I. Vorobyov, A. D. MacKerell, and R. W. Pastor. 2010. Update of the CHARMM All-Atom Additive Force Field for Lipids: Validation on Six Lipid Types. J. Phys. Chem. B 114:7830–7843.

45. Jorgensen, W. L., J. Chandrasekhar, J. D. Madura, R. W. Impey, and M. L. Klein. 1983. Comparison of simple potential functions for simulating liquid water. J. Chem. Phys. 79:926–935.

46. Neria, E., S. Fischer, and M. Karplus. 1996. Simulation of activation free energies in molecular systems. J. Chem. Phys. 105:1902.

47. Durell, S. R., B. R. Brooks, and A. Ben-Naim. 1994. Solvent-Induced Forces between Two Hydrophilic Groups. J. Phys. Chem. 98:2198–2202.

48. Beglov, D., and B. Roux. 1994. Finite Representation of an Infinite Bulk System: Solvent Boundary Potential for Computer Simulations. J. Chem. Phys. 100:9050–9063.

49. Luo, Y., and B. Roux. 2010. Simulations of Osmotic Pressure in Concentrated Aqueous Salt Solutions. J. Phys. Chem. Lett. 1:183–189.

50. Jo, S., T. Kim, and W. Im. 2007. Automated Builder and Database of Protein/Membrane Complexes for Molecular Dynamics Simulations. PLoS One 2:e880.

51. Berendsen, H. J. C., J. P. M. Postma, W. F. van Gunsteren, A. DiNola, and J. R. Haak. 1984. Molecular dynamics with coupling to an external bath. J. Chem. Phys. 81:3684–3690.

52. Hoover, W. G. 1985. Canonical dynamics: Equilibrium phase-space distributions. Phys. Rev. A 31:1695–1697.

53. Nosé, S., and M. L. Klein. 1983. Constant pressure molecular dynamics for molecular systems. Mol. Phys. 50:1055–1076.

54. Parrinello, M., and A. Rahman. 1981. Polymorphic transitions in single crystals: A new molecular dynamics method. J. Appl. Phys. (Melville, NY, U. S.) 52:7182–7190.

55. Hess, B. 2008. P-LINCS: A Parallel Linear Constraint Solver for Molecular Simulation. J. Chem. Theory Comput. 4:116–122.

56. Darden, T., D. York, and L. Pedersen. 1993. Particle mesh Ewald: An N·log(N) method for Ewald sums in large systems. J. Chem. Phys. 98:10089–10092.

57. Essmann, U., L. Perera, M. L. Berkowitz, T. Darden, H. Lee, and L. G. Pedersen. 1995. A smooth particle mesh Ewald method. J. Chem. Phys. 103:8577–8593.

58. Schmidt, T. H., and C. Kandt. 2012. LAMBADA and InflateGRO2: Efficient Membrane Alignment and Insertion of Membrane Proteins for Molecular Dynamics Simulations. J. Chem. Inf. Model. 52:2657–2669.

59. Kandt, C., W. L. Ash, and D. P. Tieleman. 2007. Setting up and running molecular dynamics simulations of membrane proteins. Methods 41:475–488.

60. Brooks, B. R., C. L. Brooks, A. D. MacKerell, L. Nilsson, R. J. Petrella, B. Roux, Y. Won, G. Archontis, C. Bartels, S. Boresch, A. Caflisch, L. Caves, Q. Cui, A. R. Dinner, M. Feig, S. Fischer, J. Gao, M. Hodoscek, W. Im, K. Kuczera, T. Lazaridis, J. Ma, V. Ovchinnikov, E. Paci, R. W. Pastor, C. B. Post, J. Z. Pu, M. Schaefer, B. Tidor, R. M. Venable, H. L. Woodcock, X. Wu, W. Yang, D. M. York, and M. Karplus. 2009. CHARMM: The Biomolecular Simulation Program. J. Comput. Chem. 30:1545–1614.

61. Schrodinger, LLC. 2015. The PyMOL Molecular Graphics System, Version 1.8.

62. Pettersen, E. F., T. D. Goddard, C. C. Huang, G. S. Couch, D. M. Greenblatt, E. C. Meng, and T. E. Ferrin. 2004. UCSF Chimera—A visualization system for exploratory research and analysis. J. Comput. Chem. 25:1605–1612.

63. Humphrey, W., A. Dalke, and K. Schulten. 1996. VMD: Visual molecular dynamics. J. Mol. Graphics 14:33–38.

64. Daura, X., K. Gademann, B. Jaun, D. Seebach, W. F. van Gunsteren, and A. E. Mark. 1999. Peptide Folding: When Simulation Meets Experiment. Angew. Chem., Int. Ed. 38:236–240.

65. Sun, Z.-Y. J., K. J. Oh, M. Kim, J. Yu, V. Brusic, L. Song, Z. Qiao, J.-h. Wang, G. Wagner, and E. L. Reinherz. HIV-1 Broadly Neutralizing Antibody Extracts Its Epitope from a Kinked gp41 Ectodomain Region on the Viral Membrane. Immunity 28:52–63.

66. Williams, L. D., G. Ofek, S. Schätzle, J. R. McDaniel, X. Lu, N. I. Nicely, L. Wu, C. S. Lougheed, T. Bradley, M. K. Louder, K. McKee, R. T. Bailer, S. O’Dell, I. S. Georgiev, M. S. Seaman, R. J. Parks, D. J. Marshall, K. Anasti, G. Yang, X. Nie, N. L. Tumba, K. Wiehe, K. Wagh, B. Korber, T. B. Kepler, S. Munir Alam, L. Morris, G. Kamanga, M. S. Cohen, M. Bonsignori, S.-M. Xia, D. C. Montefiori, G. Kelsoe, F. Gao, J. R. Mascola, M. A. Moody, K. O. Saunders, H.-X. Liao, G. D. Tomaras, G. Georgiou, and B. F. Haynes. 2017. Potent and broad HIV-neutralizing antibodies in memory B cells and plasma. Sci. Immunol. 2.

67. Yi, H. A., B. Diaz-Rohrer, P. Saminathan, and A. Jacobs. 2015. Membrane proximal external region of gp41 from HIV-1 strains HXB2 and JRFL has different sensitivity to alanine mutation. Biochemistry 54:1681–1693.

68. Owens, R. J., C. Burke, and J. K. Rose. 1994. Mutations in the membrane-spanning domain of the human immunodeficiency virus envelope glycoprotein that affect fusion activity. J. Virol. 68:570–574.

69. Murphy, R. E., A. B. Samal, J. Vlach, and J. S. Saad. 2017. Solution Structure and Membrane Interaction of the Cytoplasmic Tail of HIV-1 gp41 Protein. Structure 25:1708–1718.

70. Li, L., I. Vorobyov, and T. W. Allen. 2013. The Different Interactions of Lysine and Arginine Side Chains with Lipid Membranes. J. Phys. Chem. B 117:11906–11920.

71. Sands, Z. A., and M. S. P. Sansom. 2007. How Does a Voltage Sensor Interact with a Lipid Bilayer? Simulations of a Potassium Channel Domain. Structure 15:235–244.

72. Cleghorn, F. R., N. Jack, J. K. Carr, J. Edwards, B. Mahabir, A. Sill, C. B. McDanal, S. M. Connolly, D. Goodman, R. Q. Bennetts, T. R. O’Brien, K. J. Weinhold, C. Bartholomew, W. A. Blattner, and M. L. Greenberg. 2000. A distinctive clade B HIV type 1 is heterosexually transmitted in Trinidad and Tobago. Proc. Natl. Acad. Sci. U. S. A. 97:10532–10537.

73. Loxton, A. G., F. Treurnicht, A. Laten, E. J. van Rensburg, and S. Engelbrecht. 2005. Sequence analysis of near full-length HIV type 1 subtype D primary strains isolated in Cape Town, South Africa, from 1984 to 1986. AIDS Res. Hum. Retroviruses 21:410–413.

74. Heipertz, R. A., O. Ayemoba, E. Sanders-Buell, K. Poltavee, P. Pham, G. H. Kijak, E. Lei, M. Bose, S. Howell, A. M. O’sullivan, A. Bates, T. Cervenka, J. Kuroiwa, A. Akintunde, O. Ibezim, A. Alabi, O. Okoye, M. Manak, J. Malia, S. Peel, M. Maisaka, D. Singer, R. J. O’Connell, M. L. Robb, J. H. Kim, N. L. Michael, O. Njoku, and S. Tovanabutra. 2016. Significant contribution of subtype G to HIV-1 genetic complexity in Nigeria identified by a newly developed subtyping assay specific for subtype G and CRF02_AG. Medicine 95:e4346.

75. Vostrikov, V. V., B. A. Hall, D. V. Greathouse, R. E. Koeppe, and M. S. P. Sansom. 2010. Changes in Transmembrane Helix Alignment by Arginine Residues Revealed by Solid-State NMR Experiments and Coarse-Grained MD Simulations. JACS 132:5803–5811.

76. Freites, J. A., D. J. Tobias, G. von Heijne, and S. H. White. 2005. Interface connections of a transmembrane voltage sensor. Proc. Natl. Acad. Sci. U. S. A. 102:15059–15064.

77. Cheng, J.-X., S. Pautot, D. A. Weitz, and X. S. Xie. 2003. Ordering of water molecules between phospholipid bilayers visualized by coherent anti-Stokes Raman scattering microscopy. Proc. Natl. Acad. Sci. U. S. A. 100:9826–9830.

78. Chojnacki, J., D. Waithe, P. Carravilla, N. Huarte, S. Galiani, J. Enderlein, and C. Eggeling. 2017. Envelope glycoprotein mobility on HIV-1 particles depends on the virus maturation state. Nature Communications 8:545.

79. Gofman, Y., T. Haliloglu, and N. Ben-Tal. 2012. The Transmembrane Helix Tilt May Be Determined by the Balance Between Precession Entropy and Lipid Perturbation. Journal of Chemical Theory and Computation 8:2896–2904.

